# A miniature CRISPR-Cas10 enzyme confers immunity by an inverse signaling pathway

**DOI:** 10.1101/2025.03.28.646030

**Authors:** Erin E. Doherty, Benjamin A. Adler, Peter H. Yoon, Kendall Hsieh, Kenneth Loi, Emily G. Armbuster, Arushi Lahiri, Cydni S. Bolling, Xander E. Wilcox, Amogha Akkati, Anthony T. Iavarone, Joe Pogliano, Jennifer A. Doudna

## Abstract

Microbial and viral co-evolution has created immunity mechanisms involving oligonucleotide signaling that share mechanistic features with human anti-viral systems^1^. In these pathways, including CBASS and type III CRISPR systems in bacteria and cGAS-STING in humans, oligonucleotide synthesis occurs upon detection of virus or foreign genetic material in the cell, triggering the antiviral response^2–4^. In a surprising inversion of this process, we show here that the CRISPR-related enzyme mCpol synthesizes cyclic oligonucleotides constitutively as part of an active mechanism that maintains cell health. Cell-based experiments demonstrated that the absence or loss of mCpol-produced cyclic oligonucleotides triggers cell death, preventing spread of viruses that attempt immune evasion by depleting host cyclic nucleotides. Structural and mechanistic investigation revealed mCpol to be a di-adenylate cyclase whose product, c-di-AMP, prevents toxic oligomerization of the effector protein 2TMβ. Analysis of cells by fluorescence microscopy showed that lack of mCpol allows 2TMβ-mediated cell death due to inner membrane collapse. These findings unveil a powerful new defense strategy against virus-mediated immune suppression, expanding our understanding of oligonucleotides in cell health and disease. These results raise the possibility of similar protective roles for cyclic oligonucleotides in other organisms including humans.

## Introduction

Oligonucleotide-based immune signaling is conserved across the tree of life^1^. In all known immune signaling pathways, including prokaryotic cyclic oligonucleotide-based antiphage signaling systems (CBASS) and type III CRISPR-Cas adaptive immunity as well as eukaryotic cyclic GMP–AMP Synthase-Stimulator of Interferon Genes (cGAS-STING)^2–4^, a signaling enzyme senses viral infection and synthesizes an oligonucleotide product to activate an immune response^2,3,5–13^.

Distinct pathways for oligonucleotide-based immune signaling in bacteria have been tentatively linked by an uncharacterized gene encoding minimal CRISPR polymerase (mCpol)^14^ that often co-occurs with CBASS genes and shares sequence homology with *cas10*, the signature gene of type III CRISPR systems^15^ (Fig. 1a). These associations suggested that mCpol might contribute to bacterial immunity by an unknown mechanism. Here we show that mCpol and its associated 2TMβ effector protein confer antiphage immunity in a process that foils viral anti-immunity enzymes by inverting the role of signaling oligonucleotides. Instead of oligonucleotide-induced cell death by effector activation in response to viral infection, mCpol constitutively synthesizes 3’-5’-cyclic diadenylate (c-di-AMP) that maintains cell health by preventing toxic 2TMβ oligomerization. Disruption of c-di-AMP production in mCpol/2TMβ-containing cells, such as occurs with phage-encoded anti-CBASS (Acb) phosphodiesterases, allows 2TMβ multimerization and cell death by inner membrane collapse. These mCpol inverse signaling systems (MISS) co-locate with CBASS immune systems in bacterial genomes to establish a signaling trap in which phage anti-immunity enzymes trigger host cell destruction.

**Figure 1.**
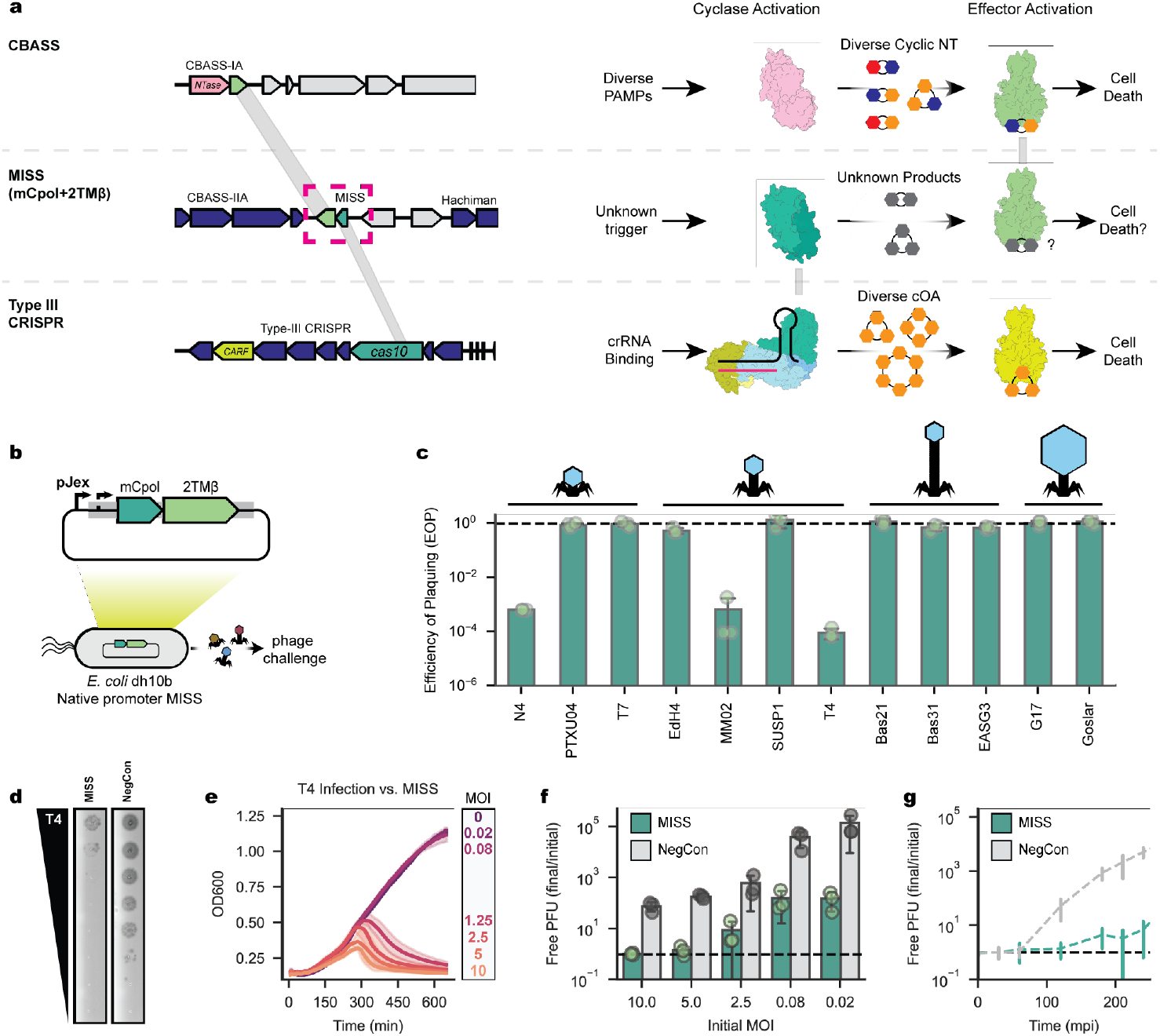
Effects of mCpol and 2TMβ expression on *E. coli* susceptibility to bacteriophage infection. **a**, Comparison of representative CBASS (top, NZ_PTRT01000132), MISS (middle, NZ_LYAF01000004)^14^ and Type III-A CRISPR-Cas loci (bottom, NC_002976) shows shared protein domains between CBASS and MISS (2TMβ and Cap15, 23% sequence identity) and between MISS and Type III CRISPR loci (mCpol and Cas10, 19% sequence identity). Shared protein domains are highlighted. CBASS, MISS and Type III CRISPR cyclic nucleotide signaling architecture in response to pathogen associated molecular patterns (PAMPs) is shown on the right. **b**, MISS expression for phage challenge experiments. MISS locus from ECOR31 is expressed on a low copy plasmid under control of its native promoter. **c**, Change in efficiency of plaquing when phages of 12 unique phage genera were challenged against *E. coli* with ECOR31-derived MISS loci normalized to a vector control (Methods). Data shown represent the mean of three biological replicates ± standard deviation. The black dashed line represents no change in EOP. **d**, Representative plaque assay for ECOR31-derived MISS versus a negative control (NegCon) against phage T4. Assay shown is representative of three biological replicates. **e**, MISS-dependent protection against phage T4 shows incomplete protection at high MOI, but population-level protection at low MOI. Data shown represent the mean of three biological replications ± standard deviation. **f**, T4 phage production assay following overnight phage infection for MISS and a vector control. Data are presented as the mean of 3 biological replicates ± standard deviation. The black dashed line represents no additional production of phages compared to the beginning of the experiment. **g**, One-step growth curve of T4 infection MISS and a vector control, representing one round of infection at ∼0.01 MOI. Data are presented as the mean of three biological replicates ± standard deviation. The black dashed line represents no additional production of phages.

## Results

### mCpol contributes to antiphage defense

While investigating the genomic locus surrounding the Hachiman antiphage defense system in *E. coli* ECOR31^16^, we observed an adjacent operon encoding mCpol and a 2TMβ protein, hereon referred to as MISS (Fig. 1a). To test whether mCpol plays an independent role in antiphage immunity, we expressed the MISS operon, containing just the mCpol and 2TMβ genes, using its native promoter in *E. coli* and challenged the strain with 12 different diverse bacteriophages (Fig. 1b). Although most phage killed the cells in the presence of MISS, expression of MISS provided 10^3^-10^4^-fold protection from phages T4, MM02 and N4 (Fig. 1c, d; Supplementary Fig. 1). To determine the mode of protection, we exposed the MISS-expressing strain to phage T4 at various multiplicities of infection (MOIs). High MOIs (≥1.25 infections per cell) resulted in limited phage production but also significant cell death (Fig. 1e, f). At low MOI (≤0.08 infections per cell), however, MISS limited T4 production in early rounds of infection to provide population-level anti-phage protection (Fig. 1g). Together, these results suggest that MISS confers phage defense through an abortive infection mechanism consistent with its encoding of a 2TMβ effector homolog^17^.

Having established MISS as an antiphage system, we next investigated the basis of MISS signaling. An alignment of 163 mCpol proteins revealed a semi-conserved di-glycine diaspartate (GGDD) sequence similar to the active-site residues in Cas10 and related nucleotidyl cyclases that catalyze ATP dicyclization (Supplementary Fig. 2)^15,18,19^. To assess whether mCpol has the intrinsic ability to synthesize cyclic oligonucleotides, recombinant mCpol was incubated in a buffer containing adenosine or guanosine triphosphate (ATP or GTP). Analysis of these reactions by thin layer chromatography (TLC) showed the presence of new species in the ATP-containing reactions but not those containing GTP (Fig. 2a). Mutation of the strictly conserved aspartate residue (D57A) in mCpol completely eliminated production of one new species (Fig. 2a; Supplementary Fig. 2). Treatment of mCpol-ATP reaction products with S1 nuclease, which specifically cleaves 5’-3’ phosphodiester linkages, also depleted the reaction of the D57-dependent product (Fig. 2a).

**Figure 2.**
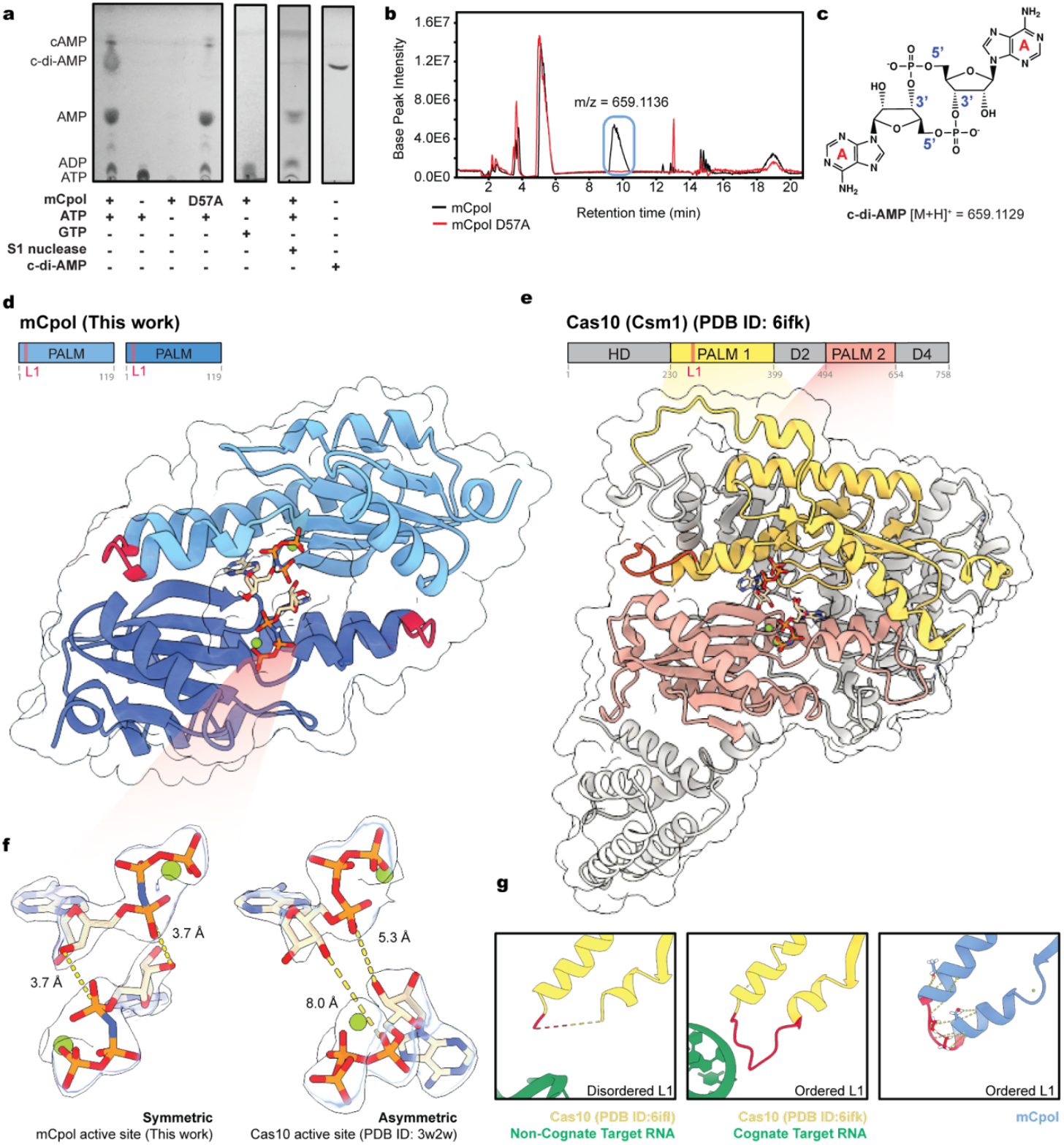
Crystal structure of mCpol and biochemical detection of adenylate cyclase activity. **a**, TLC analysis of ATP-specific product formation upon reaction with mCpol. The primary product of mCpol reaction is degraded by S1 nuclease and co-migrates with c-di-AMP. **b**, Differential HPLC-MS reveals c-di-AMP as the primary product of mCpol. **c**, Structure and molecular weight of primary mCpol product, c-di-AMP. **d**, X-ray crystal structure of mCpol bound to ApNHpp. **e**, Domain map and structure of Cas10 (Csm1) highlighting Palm domains conserved between mCpol and Cas10 (PDB ID: 6ifk). **f**, Positioning of 2 ATP analogs in mCpol active site (this work) and 2 ATP in Csm1 active site (donor/acceptor) (PDB ID: 3w2w). **g**, The L1 loop of Csm1 is disordered in the presence of non-cognate target RNA (PDB ID: 6ifl) and ordered in the presence of cognate target RNA (PDB ID: 6ifk). The region of mCpol corresponding to Loop L1 consists of a highly ordered turn stabilized by intramolecular hydrogen bonds.

To determine the identity of mCpol’s primary reaction product, we used differential high-performance liquid chromatography-mass spectrometry (HPLC-MS) to show that the single species present with wild-type mCpol but absent with the D57A mutant is 5’-3’ cyclic di-adenosine monophosphate (c-di-AMP) (Fig. 2b, c). TLC analysis using a chemical standard showed that the mCpol-ATP reaction product co-migrates with c-di-AMP, and TLC-MS confirmed the product’s corresponding molecular weight (Fig. 2a; Supplementary Fig. 3). These results reveal that mCpol synthesizes c-di-AMP from ATP, mirroring the selectivity of Cas10 for the adenine base^8^. Notably, however, mCpol makes a singular c-di-AMP product whereas Cas10 makes an array of higher order cyclic oligoadenylates (cOA_n_; n = 2 to 6)^7^.

**Figure 3.**
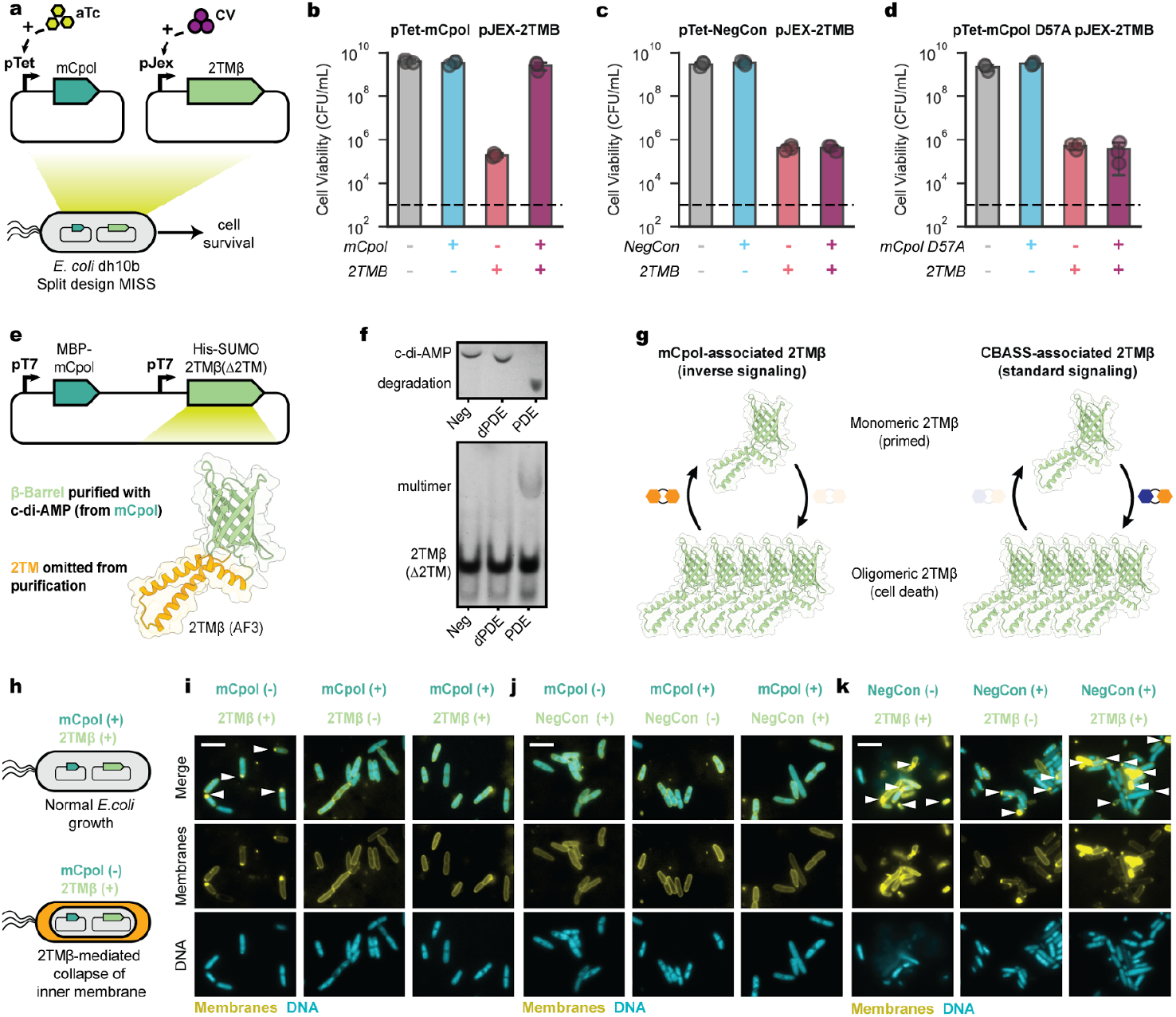
Effects of mCpol-generated c-di-AMP on 2TMβ protein behavior. **a**, Overview of mCpol split-design to investigate toxin-antitoxin dynamics. For toxin-antitoxin assays in b-d, toxin and antitoxin components are expressed under a pJex promoter with 0 or 50 nM crystal violet (CV) inducer and under a pTet promoter with 0 or 20 nM anhydrotetracycline inducer (aTc) respectively. **b**, 2TMβ-induced reduction in CFU (colony forming units) is subverted by mCpol expression. **c**, 2TMβ reduces cell viability in the absence of mCpol. **d**, Toxicity of 2TMβ is not suppressed by catalytically deactivated mCpol (D57A). **e**, Approach for purification of 2TMβ (Δ2TM) in the presence of mCpol. Model generated by AlphaFold 3 (AF3). **f**, TLC showing degradation of c-di-AMP by a phosphodiesterase (PDE) and lack of degradation by a catalytically inactivated PDE (dPDE) or no enzyme (Neg). 2TMβ multimerizes upon depletion of c-di-AMP by a PDE^42^, but not in the presence of dPDE. **g**, Contrasting mechanisms of MISS-associated 2TMβ (left) and CBASS-associated 2TMβ (right). CBASS-associated 2TMβ mechanism is adapted from Duncan-Lowey et al^17^. **h**, Bacterial physiology of inner membrane integrity in the presence (top) or absence (bottom) of mCpol expression. For microscopy experiments shown in panels i-j, mCpol and candidate toxin are expressed by induction with 0 or 200 nM aTc and 0 or 50 nM respectively. DNA is stained with 4’,6-diamidino-2-phenylindole (DAPI, teal) and all membranes are stained with MitoTracker Green (yellow). Scale bar represents 5 µm. Arrows highlight polar membrane collapse or lack thereof. **i**, Fluorescence microscopy shows 2TMβ-mediated formation of membrane lesions (white arrows) if expressed in the absence of mCpol. **j**, Fluorescence microscopy shows no loss of inner membrane integrity for a 2TMβ vector control. **k**, Fluorescence microscopy shows loss of inner membrane integrity for leaky (−) and induced (+) expression of 2TMβ in a strain with a vector control lacking mCpol.

### mCpol-generated c-di-AMP signaling maintains cell health

Previously studied antiviral second messenger synthases are tightly regulated sensors that become active during infection^20^. For example, cOA synthesis by Cas10 depends on cognate target RNA recognition^21^. However, we observed that c-di-AMP synthesis by mCpol proceeded *in vitro* without an additional stimulus. To determine how mCpol’s regulation may diverge from that of Cas10, we solved a high resolution X-ray crystal structure of mCpol bound to a non-hydrolyzable ATP analog (ApNHpp) (Fig. 2d; Supplementary Fig. 4). This structure showed mCpol to contain a Cas10-like Palm domain that dimerizes to form a pocket where two ATP analogs are bound (Fig. 2d, e)^22^. Conserved serine residues, also found in Cas10, confer ATP binding selectivity (Supplementary Fig. 5). However, the positioning of ATP substrates in the active site differs substantially between mCpol and Cas10. Previous structures of Cas10 bound to ATP revealed the presence of asymmetric donor and acceptor molecules positioned to allow for a stepwise formation of each 5’ to 3’ linkage (Fig. 2f)^21,23^. In the mCpol active site, however, symmetry between the two ATP analogs positions them for concurrent synthesis of both 5’-3’ linkages to form c-di-AMP (Fig. 2f). This substrate orientation explains the singular product identity that we observed biochemically.

**Figure 4.**
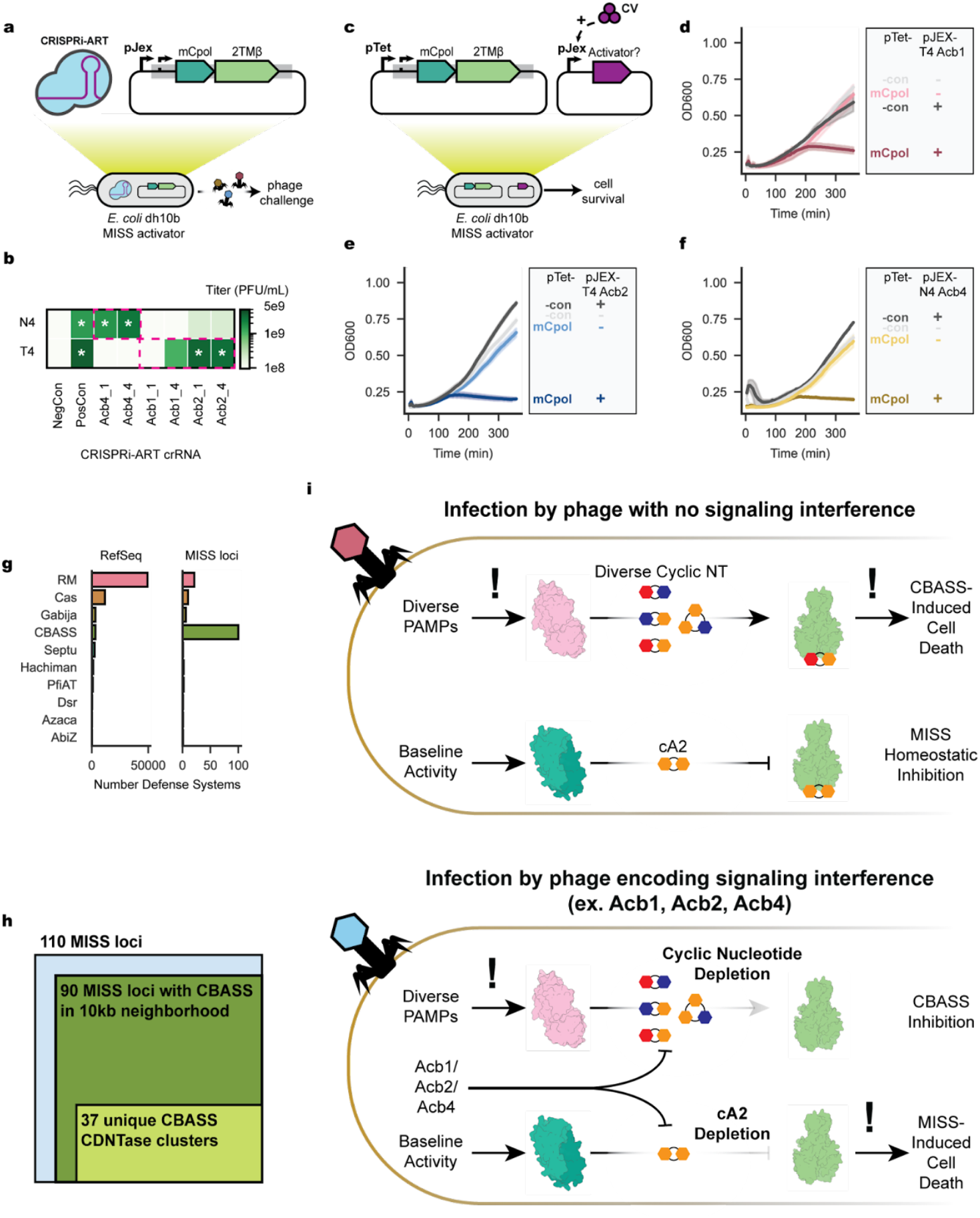
MISS activation by phage encoded anti-CBASS (Acb) proteins. **a**, CRISPRi-ART screening of phage encoded Acb proteins to determine phage infectivity in the context of MISS. **b**, Heatmap of CRISPRi-ART screening of phage encoded Acb proteins during T4 or N4 infection colored by plaque forming units (PFU/mL). For all experiments, CRISPRi-ART was expressed at 20 nM anhydrotetracycline inducer (aTc). Negative control (NegCon) represents a CRISPRi-ART crRNA targeting a transcript not present in the experiment. Positive control (PosCon) represents a non-targeting CRISPRi-ART crRNA in the absence of MISS. Restoration of plaques to PosCon size are marked with a white asterisk. CRISPRi-ART conditions targeting a phage-encoded Acb protein in the target phage are bounded in dashed magenta lines. Phage PFU/mL are reported as the mean of 3 biological replicates. **c**, Overview of activator assays in the context of MISS under control of its native promoter. For activator assays in panels d-f candidate MISS activator proteins are expressed at 0 or 125 nM crystal violet (CV). **d**, T4 Acb1 expression is sufficient to activate MISS toxicity. **e**, T4 Acb2 expression is sufficient to activate MISS toxicity. **f**, N4 Acb4 expression is sufficient to activate MISS toxicity. **g**, DefenseFinder quantification of all known defense systems as of 2024 (left) versus the 10kb neighborhood of mCpol-containing proteins. **h**, Fraction of CBASS-containing loci nearby MISS systems. **i**, Schematic of proposed evolutionary trap between CBASS and MISS_TA defense systems. During normal phage infection (top), MISS_TA signaling is uninterrupted and MISS_TA defense is not activated. However, the phage could be susceptible to CBASS-mediated defense. During infection by phages that broadly interfere with cyclic nucleotide signaling (ex. T4, MM02, N4), the phage can escape CBASS defense through cyclic nucleotide depletion or sequestration. However, c-di-AMP is also depleted, activating MISS_TA defense.

Previous studies have shown that Loop L1 of the Cas10 Palm 1 domain (*Sth*Csm1 residues 257-265) acts as a critical allosteric regulator of cyclase activity^24^. This regulatory loop bridges two alpha helices, one of which borders the active site residues. Residues in Cas10 Loop L1 are disordered in the presence of non-cognate target RNA, which does not activate Cas10, but become ordered upon binding of cognate target RNA (Fig. 2g)^24^. In the structure of mCpol, however, a highly ordered turn that is stabilized by a network of hydrogen bonds replaces the region corresponding to Cas10’s Loop L1 (residues 19-23). This structural difference suggests that mCpol adopts an active conformation without additional requirements such as RNA binding (Fig. 2g). The absence of homology to Cas10’s regulatory region^21,24^, and the stabilization of the Loop L1-equivalent structure, are consistent with our observation that mCpol is a constitutively active cyclase.

We wondered how constitutive c-di-AMP synthesis enables an immune response specific to phage infection. Early in our investigation of mCpol-mediated phage defense, we were surprised to observe that increased expression of mCpol-2TMβ conferred reduced phage defense (Supplementary Fig. 6), since increased expression of the CBASS nucleotidyltransferase and effector has the opposite effect of providing more robust phage defense and background toxicity^17^. Based on our observations of constitutive mCpol activity and increased mCpol expression leading to decreased phage defense and cell death, we hypothesized that the MISS operon acts by mCpol-mediated negative regulation, akin to toxin-antitoxin systems.

To test for a toxin-antitoxin relationship between mCpol and 2TMβ, we separated the MISS operon into two plasmids, where orthogonal inducers controlled expression of mCpol and 2TMβ (Fig. 3a). In the absence of mCpol, we found that 2TMβ expression had a deleterious effect on cell viability and growth, suggesting its role as a toxin (Fig. 3c). mCpol expression suppressed the growth defects incurred by 2TMβ expression, suggesting that it plays a role in neutralizing 2TMβ (Fig. 3b). No suppression of 2TMβ toxicity occurred upon coexpression of an mCpol catalytic mutant (Fig. 3d). These data indicate that MISS is a toxin-antitoxin system in which the catalytic activity of mCpol prevents 2TMβ toxicity.

### c-di-AMP blocks 2TMβ-induced cell lysis

The observation that active mCpol suppresses toxicity of 2TMβ suggested that mCpol’s c-di-AMP product operates as the cognate antitoxin. 2TMβ comprises N-terminal transmembrane helices and a C-terminal β-barrel domain in an architecture mirroring the CBASS effector protein Cap15 (Fig. 3e). Upon cyclic dinucleotide binding to its β-barrel domain, Cap15 multimerizes and induces cell death by disrupting inner-membrane integrity^17^. We noted that residues involved in cyclic oligonucleotide binding by Cap15 are conserved in 2TMβ (Supplementary Fig. 7), suggesting that c-di-AMP binding may directly control 2TMβ activity. To test the hypothesis that mCpol-generated c-di-AMP inhibits 2TMβ multimerization-mediated cell death, we examined the effect of c-di-AMP depletion on purified 2TMβ. The β-barrel domain of 2TMβ (2TMβ Δ2TM) was overexpressed in the presence of mCpol and purified with supplementation of c-di-AMP in all buffers (Fig. 3e). When a 2H-phosphodiesterase (PDE) was used to degrade c-di-AMP in the sample containing recombinant 2TMβ Δ2TM, we observed a shift of the protein to a higher molecular weight species (Fig. 3f). However, when catalytically inactivated PDE (H>A, H>R; dPDE) was used, the shift was abrogated (Fig. 3f). These results suggest that, rather than multimerizing in response to the cyclic dinucleotide like the CBASS Cap15 effector, 2TMβ multimerizes in the absence of c-di-AMP (Fig. 3g).

We next analyzed the basis for 2TMβ’s toxicity by assessing bacterial cell integrity under different conditions using fluorescence microscopy (Fig. 3h-j). When 2TMβ expression alone was induced in our co-expression strain (Fig. 3i), cells developed membrane lesions, most often at the cell poles. Consistent with our cell viability assays (Fig. 3b-c), membrane lesions did not develop when mCpol alone or both mCpol and 2TMβ were expressed (Fig. 3i) or in our negative control strain carrying an empty vector instead of the 2TMβ plasmid (Fig. 3j). Cells lacking the mCpol plasmid (2TMβ only) developed membrane lesions and frequently lysed (Fig. 3k). Collectively, these results demonstrate that mCpol signaling prevents cell death by inhibiting 2TMβ multimerization-mediated inner membrane collapse. This inverse regulatory logic deviates from all known immune signaling mechanisms in which virus-triggered nucleotide signals activate an immune effector.

### Acb proteins activate 2TMβ toxicity

The inverse signaling logic encoded by MISS suggested that disruption of immune signaling could trigger 2TMβ-induced cell death. Notably, phages sensitive to MISS encode Acb proteins that interfere with immune signaling by hydrolyzing or sequestering cyclic oligonucleotides (Fig. 1c). T4 and MM02 harbor cyclic nucleotide degrader Acb1 (encoded by *T457B* and MM02*gp156*)^25^ and cyclic nucleotide sponge Acb2 (encoded by *T4vs*.*4* and *MM02gp117*)^26^. N4 harbors cyclic nucleotide sponge Acb4 (*N4gp48*)^27^. Furthermore, three T4-like phages that are naturally resistant to MISS encode loss-of-function mutations in *acb1* or *acb2* (Supplementary Fig. 8)^28,29^. To test whether Acb1_T4_, Acb2_T4_ and Acb4_N4_ are necessary for MISS activation, we used CRISPRi-ART^30^ to disrupt expression of these phage genes during infection (Fig. 4a; Supplementary Fig. 9). CRISPRi-ART elimination of Acb proteins (Acb4_N4_ and Acb2_T4_) restored infectivity to levels similar to those observed in the absence of MISS (Fig. 4b). These results suggest that, in the context of phage infection, Acb expression is necessary for activating MISS defense.

To test whether Acb proteins are sufficient to activate MISS in the absence of phage infection, we coexpressed MISS and candidate activator proteins Acb1_T4_, Acb2_T4_ and Acb4_N4_^25–27^ (Fig. 4c). Strikingly, we found that low levels of Acb1_T4_ expression were sufficient to activate MISS-induced cell death (Fig. 4d), while an enzymatically inactivated Acb1_T4_ was insufficient (Supplementary Fig. 10). Similarly, low levels of Acb2_T4_ expression were sufficient to activate MISS-induced cell death (Fig. 4e), while a known Acb2_T4_ binding site mutant^26^ was not (Supplementary Fig. 10). The cyclic nucleotide sponge protein Acb_N4_^27^ was also sufficient to activate MISS-induced cell death (Fig. 4f). Collectively, these results demonstrate that Acb signaling interference proteins are necessary and sufficient to activate an immune response from MISS.

Antiphage systems tend to co-localize alongside other defense-associated operons in genetic islands^31,32^ and can exhibit synergistic activity through either complementary phage resistance patterns or mechanistic traps^33,34^. Given that diverse Acb proteins were both necessary and sufficient to activate MISS-induced cell death, we suspected that CBASS systems would be enriched near mCpol-containing genomic loci. To explore this possibility, we analyzed all publicly available mCpol-encoding genes and their genomic neighborhoods, identifying 295 unique mCpol-encoding loci. Phylogenetic analysis of *mCpol* genes highlighted a clade of 109 unique mCpol-encoding loci representing MISS (Supplementary Fig. 11, blue highlight). DefenseFinder^35^ analysis of these 109 gene neighborhoods identified 91 unique genes encoding CBASS nucleotidyltransferases across 89 of these loci, reflecting a strong enrichment of CBASS relative to the general distribution of defense systems in bacteria^35^ (Fig. 4g, h). These CBASS nucleotidyltransferase genes reflected at least 37 unique clusters (80% coverage, 40% identity), suggesting many distinct events of co-association between MISS and CBASS (Fig. 4i; examples shown in Supplementary Fig. 11b). We propose that the strong co-occurrence between CBASS and MISS reflects an immune signaling trap (Fig. 4i). Through inverse immune signaling alongside CBASS, MISS limits the spread of phages that interfere with CBASS immune signaling.

## Discussion

Our results reveal that the MISS locus encodes a defense system composed of a Cas10-like oligonucleotide cyclase (mCpol) and a CBASS-like effector protein (2TMβ) that counters phage-encoded anti-defense mechanisms. mCpol is a constitutively active enzyme that produces c-di-AMP to homeostatically inhibit oligomerization of its 2TMβ effector. The loss of c-di-AMP triggers 2TMβ oligomerization and cell death due to inner membrane collapse, a process activated by phage anti-defense proteins, such as anti-CBASS, which deplete cyclic oligonucleotide messengers^25–27,36–39^. When triggered early in the phage infection process, infection aborts due to cell death that protects other cells in a microbial population. These results extend discoveries of virus-encoded anti-immunity proteins^25–27,36–39^ to show how host systems can counter virus-mediated immune suppression.

MISS is the first known immune signaling system to exhibit a toxin-antitoxin mechanism of regulation in which the signaling molecule maintains cell health through effector suppression. There are currently eight described RNA or protein antitoxin types, where classification is based on the antitoxin mechanism^41^. Given that mCpol-produced c-di-AMP serves as the antitoxin to the cognate 2TMβ toxin, we propose that MISS represents a novel class of TA system (Type IX) in which the antitoxin is a small molecule produced by a dedicated antitoxin gene.

Mechanisms of signaling immunity and immune evasion are conserved across kingdoms of life, as evidenced by homology between enzymes responsible for cGAS-like signaling in prokaryotes and eukaryotes^3,9^ in which cyclic oligonucleotides induce an immune response that limits viral propagation. Correspondingly, 2H-phosphodiesterases encoded by eukaryotic and prokaryotic viruses deplete these signals to evade immune detection^42,43^. Therefore, these immune systems are subject to cross-kingdom evolutionary pressures which likely gave rise to the inverse signaling mechanism seen in MISS. The discovery of a Type IX toxin-antitoxin mechanism in prokaryotes suggests that inverse signaling architectures in eukaryotic immunity are likely to be uncovered.

MISS counters phage anti-defense proteins that promiscuously disrupt cyclic oligonucleotides, acting as a safeguard of immune signaling pathways in antiviral defense. Informatic analysis of closely related MISS loci suggests a specific, mechanistic synergy with diverse CBASS systems^33^. This proposed synergy between MISS and CBASS immune signaling pathways implies a selective pressure for anti-immune proteins to develop more precise recognition of immune signaling molecules, which likely comes at the cost of broad-spectrum immune evasion. Collectively, our results reveal a major constraint for the evolution of viral proteins that inhibit signaling immune systems and suggest the possibility of similar protective roles for cyclic oligonucleotides across the tree of life.

## Acknowledgements

We thank members of the Doudna lab and collaborators including Dr. David Colognori, Dr. Jason Nomburg, Owen Tuck, Dr. Daniel Belieny, Dr. Linxing Chen, Dr. Isabel Esain-Garcia and Dr. Santiago Lopez for discussion, encouragement, and feedback. J.A.D. is an investigator of the Howard Hughes Medical Institute, and research in the Doudna laboratory is supported by the Howard Hughes Medical Institute (HHMI), NIH/NIAID (U19AI171110, U54AI170792, U19AI135990, UH3AI150552 and U01AI142817), NIH/NINDS (U19NS132303), NIH/NHLBI (R21HL173710), NSF (2334028), DOE (DE-AC02-05CH11231, 2553571 and B656358), Lawrence Livermore National Laboratory, Apple Tree Partners (24180), Mr. Li Ka Shing, Koret-Berkeley-TAU, Emerson Collective and the Innovative Genomics Institute (IGI).The authors acknowledge funding from the HHMI Emerging Pathogens Initiative grant (to J.P.) and NIH grant R01-GM129245 (to J.P.). E.E.D. was supported by NIGMS of the NIH under award number F32GM153031. B.A.A. was supported by m-CAFEs Microbial Community Analysis and Functional Evaluation in Soils, a Science Focus Area led by Lawrence Berkeley National Laboratory based on work supported by the US Department of Energy, Office of Science, Office of Biological and Environmental Research under contract number DE-AC02-05CH11231. E.G.A. was supported by an NIH PiBS training grant (T32 grant GM133351). C.S.B was supported by a University of California Office of the President funded UC-Historically Black Colleges and Universities Initiative (UC-HBC) award to the Doudna lab. X.E.W was supported by NIAID of the NIH under award number T32AI145821. Use of the Stanford Synchrotron Radiation Lightsource, SLAC National Accelerator Laboratory, is supported by the U.S. Department of Energy, Office of Science, Office of Basic Energy Sciences under Contract No. DE-AC02-76SF00515. The SSRL Structural Molecular Biology Program is supported by the DOE Office of Biological and Environmental Research, and by the National Institutes of Health, National Institute of General Medical Sciences (P30GM133894). The QB3/Chemistry Mass Spectrometry Facility received National Institutes of Health support (grant number 1S10OD020062-01). The contents of this publication are solely the responsibility of the authors and do not necessarily represent the official views of NIGMS or NIH.

## Author contributions

E.E.D, B.A.A, and J.A.D conceived of the project. E.E.D, B.A.A., K.H., E.G.A, A.L., C.S.B, and A.A. performed experiments. B.A.A, P.H.Y., and K.L. performed bioinformatics experiments and analyses. E.A.G performed microscopy experiments. E.E.D. and X.E.W. processed and refined crystallographic data. A.T.I. collected mass spectrometry data. J.P. supervised microscopy experiments. E.E.D., B.A.A., and J.A.D. wrote the original draft of the manuscript. All authors edited the manuscript and support its conclusions.

## Data availability

Plasmids will be made available through Addgene upon peer review of this study. The structure of mCpol-ApNHpp has been deposited under PDB ID: 9NWN and will be released upon publication of the primary citation.

## Materials and Methods

### Bacterial strains and growth conditions

Standard bacterial cultures were grown in lysogeny broth (LB Lennox) at 37°C and 250 rpm except where stated otherwise. When necessary, 34 µg/mL chloramphenicol (+Ch), + 50 µg/mL kanamycin sulfate (+K) or +50µg/mL carbenicillin (+C) was supplemented to media and LB agar plates. For cellular assays, all bacterial strains were stored at -80°C in 25% glycerol (Sigma) when not in use. Cloning and cellular assays were generally performed in *E. coli* DH10b genotype cells (*F – mcrA Δ(mrr-hsdRMS-mcrBC) endA1 recA1 ϕ80dlacZΔM15 ΔlacX74 araD139 Δ(ara, leu)7697 galU galK rpsL (StrR) nupG λ-*) (Intact Genomics, NEB). For phage experiments involving G17 or Goslar, *E. coli* strain ECOR47 was used.

### Phage propagation and scaling

Phages were propagated using standard protocols. Typically, phage production was performed at 37°C in LB Lennox media infecting *E. coli* BW25113 (F- DE(araD-araB)567 lacZ4787(del)::rrnB-3 LAM- rph-1 DE(rhaD-rhaB)568 hsdR514) at an initial MOI of 0.1. Phages G17 and Goslar were propagated on *E. coli* ECOR47. Phage titers were determined on their propagation hosts. Bacteriophages used in this study are listed in Supplementary File 2.

### Plasmid construction

Plasmids were assembled using PCR, gel extraction (Zymo) and Golden Gate assembly^44^ or Gibson Assembly using 25-30 bp of overlapping homology^45^. For some assemblies, Golden Gate ready DNA was ordered from TWIST BioSciences in lieu of generation by PCR. Native promoter MISS includes 300bp upstream of the gene encoding mCpol and 150bp downstream of the gene encoding 2TMβ. In general, plasmids were propagated in DH10b genotype *E. coli* (Intact Genomics). For G17 and Goslar assays, select plasmids were transferred to *E. coli* ECOR47 through electroporation. For protein expression and purification, plasmids were transferred into BL21 AI genotype *E. coli* (F- ompT hsdSB (rB-mB-) gal dcm araB::T7RNAPtetA). For assays involving two plasmids, plasmids were first cloned individually and co-transformed into DH10b genotype *E. coli* (Intact Genomics) through electroporation. All plasmids and co transformations used in this study were sequenced-confirmed by full-plasmid sequencing (Plasmidsaurus). To verify co-transformed plasmids, raw reads were mapped against reference plasmid sequences using Geneious. Plasmids used in this study are listed in Supplementary File 1.

### Phage infection assays

Phage infection plaque assays were performed via double agar overlay. Overlays were formed by mixing 100µL of saturated overnight cultures grown at 37°C, 250 rpm with 5 mL molten LB Lennox agar (0.7% w/v, 60°C). For G17 and Goslar assays, a lower percentage top agar concentration was used (0.35% w/v). The agar-bacterial mixture was supplemented with kanamycin to a final overlay concentration of 50 µg/mL. For plaque assays involving crystal violet (CV) (Sigma), crystal violet was supplemented to a final overlay concentration as specified. The top agar and bacterial mixture was poured onto a 5 mL LB Agar and K plate and left to dry under microbiological flame for 15 min. Phages were diluted tenfold in SM buffer (Teknova) and 2 µL of each dilution were spotted onto the top agar overlay and dried under microbiological flame. Once dry, plates were incubated at 30°C for 12-16 hours. Plates were scanned in a standard photo scanner (Epson) and plaque forming units (p.f.u) were enumerated, keeping note of changes in plaque size relative to a negative control. “Lysis from without” ^46^ phenotypes were interpreted as a lack of productive phage infection and approximated as 1 plaque forming unit (p.f.u.). Efficiency of plaquing calculations were calculated as mean(p.f.u in condition)/mean(p.f.u. negative control). The negative control is defined as a strain harboring a plasmid encoding red fluorescent protein (RFP) instead of MISS (pBA1326). All plaque assays were performed in biological triplicate from independent overnight cultures. Visualizations were performed using Seaborn in Python.

For plaque assays involving CRISPRi-ART^30^, experiments were performed with the following modifications. Kanamycin, chloramphenicol and anhydrotetracycline (aTc) (Sigma) were additionally added to final overlay concentrations of 50 µg/mL, 34 µg/mL and 20 nM respectively and plated onto a 5 mL LB Agar, K and Ch plate. The negative control is defined as MISS (pBA1751) alongside a CRISPRi-ART vector with an RFP-targeting guide RNA (pBA635) ^30^. The positive control is defined as a pJEX vector without MISS (pBA1326) alongside a CRISPRi-ART guide RNA targeting a phage not present in the experiment. Titers were calculated as mean (p.f.u. condition). All plaque assays were performed in biological triplicate from independent overnight cultures. Visualizations were performed using matplotlib and Seaborn in Python. CRISPRi-ART guide RNAs were chosen using “gRNA1” and “gRNA4” designs as described previously ^30^.

### Bacteriophage liquid growth assays and phage production estimation

Liquid phage experiments were performed in a Biotek plate reader using LB Lennox + K media. Strains containing native context MISS (pBA1751) or a negative control (pBA1326) plasmid were grown overnight at 37°C and 250 rpm. 8e6 c.f.u. of overnight culture were seeded into each well of a 96-well plate (Corning 3903) in 200 µL media. For phage experiments, T4 was diluted in LB+K media to achieve defined MOIs during infection except for MOI=0 in which no phage was added. Growth was monitored in a Biotek Cytation 5 plate reader for 12 h at 800 rpm shaking at 30°C with OD600 readings every 5 min. At the end of the experiment, cultures were pelleted and the supernatant from investigated wells was collected. Phage production was estimated via plaque assay (above) on *E. coli* harboring a negative control (pBA1326) plasmid and dividing by the effective titer at time 0. All liquid phage experiments were performed in biological triplicate and replicate conditions from independent overnights. Visualizations were performed using matplotlib and Seaborn in Python.

Estimation of free phage particle production from a single round of infection was performed in 5mL cultures. Cultures were inoculated with 2e8 c.f.u. *E. coli* harboring native context MISS (pBA1751) or a negative control (pBA1326) plasmid in LB + K media and incubated at 30°C, 250 rpm. for 15 min. Following incubation, ∼2e6 c.f.u. of phage T4 was added for a low MOI infection of ∼0.01. Infections proceeded at 30°C, 250 rpm and 200 µL sampled every 30 min for 6.5 h. For each 200 µL sample, remaining cells were pelleted, 100 µL supernatant was extracted and stored on ice until every sample was collected. Phage titers were enumerated via plaque assay on *E. coli* harboring a negative control plasmid (pBA1326). Free phage production was calculated by dividing the sample titer by the number of added phages. Replicates were performed in biological triplicate, sourcing samples from parallel cultures seeded from independent overnights. Visualizations were performed using matplotlib and Seaborn in Python.

### MISS toxin-antitoxin assays

To test for a potential toxin-antitoxin relationship in MISS, we cloned candidate antitoxin mCpol or mCpol D57A under control of the pTet promoter in a p15a-CmR plasmid and candidate toxin 2TMβ under control of the pJex promoter ^47^ in a low copy SC101-KanR plasmid. dCas13d under pTet control on a p15a-CmR plasmid (pBA635)^30^ was employed as an antitoxin negative control. RFP expressed under pJex control on a SC101-KanR plasmid was employed as a toxin negative control. Both sets of plasmids were co-transformed into DH10b *E. coli* (Intact Genomics) and sequence-verified using whole plasmid sequencing (Plasmidsaurus), followed by read alignment (Geneious). Sequence-confirmed cotransformants were stored at -80°C in 20% glycerol (v/v) until further use.

To perform solid agar toxin-antitoxin assays, LB Agar plates were freshly poured and dried under flame with the following supplements: 35 µg/mL chloramphenicol, 50 µg/mL kanamycin, variable amounts of crystal violet (CV) to induce toxin expression and variable amounts of aTc to induce antitoxin expression. For toxin expression conditions, +50 nM CV was added. For antitoxin expression conditions, +2 nM aTc was added. Once the agar was dried, 3 independent overnight cultures containing candidate toxin and antitoxin plasmids were plated in 10x serial dilutions with 5 µL spots and let dry under flame. Once dried, plates were incubated at 30°C overnight. To let colonies mature for imaging, plates were transferred to a 37°C incubator for an additional 24 h. Colonies were imaged and counted.

To perform liquid culture toxin-antitoxin assays, 3 independent overnight cultures containing candidate toxin and antitoxin plasmids were inoculated in LB media at 8e6 c.f.u. in a Corning 3903 plate with the following supplements: 35 µg/mL chloramphenicol, 50 µg/mL kanamycin, variable amounts of CV to induce toxin expression and variable amounts of aTc to induce antitoxin expression. For toxin expression conditions, +125 nM CV was used. For antitoxin expression conditions, +20 nM aTc was used. The plate was monitored in a Cytation5 plate reader (Biotek) at 30°C, 807 rpm and OD600 measured every 5 min for 12 h. Data were plotted using the Seaborn package in Python.

### MISS activator assays

To test for an activator relationship with MISS, we cloned MISS in its native context into a p15a-CmR plasmid and candidate activators T4Acb1, T4Acb1(H44A,H113A), T4Acb2, T4Acb2(Y8A) and N4Acb4 under control of the pJex promoter ^47^ in a low copy SC101-KanR plasmid. dCas13d under pTet on a p15a-CmR plasmid (pBA635) ^30^ was employed as MISS negative control. RFP under pJex on a SC101-KanR plasmid was employed as an activator negative control.

To perform liquid culture activator assays, 3 independent overnight cultures containing candidate MISS and candidate activator plasmids were inoculated in LB media at 8e6 CFU in a Corning 3903 plate with the following supplements: 35 µg/mL Chloramphenicol, 50 µg/mL Kanamycin, variable amounts of CV to induce candidate activator expression and no aTc for MISS expression. For candidate activator expression conditions, +125 nM CV was used. The plate was monitored in a Cytation5 plate reader (Biotek) at 30°C, 807 rpm and OD600 measured every 5 min for 12 h. Data were plotted using the Seaborn package in Python.

### Protein expression and purification

Expression sequences for ECOR31 mCpol were cloned into a custom pET-based vector by Gibson assembly to yield an N-terminal His_10_-MBP-TEV or C-terminal TEV-MBP-His_10_ construct. Expression plasmids for wild-type and H72A/H167R mutant pigeon pox virus PDEs were cloned into a custom pET-based vector by Gibson assembly to yield an N-terminal His_10_-MBP-TEV construct. Expression plasmids for ECOR31 TM2β residues 72-200 were cloned into a pET Duet-1 vector for co-expression of His_6_-SUMO2-TM2β and MBP-TEV-mCpol constructs. For studies using GFP tagged TM2β, fluorescent tags were added to the pET Duet-1 vector for the production of His_6_-SUMO2–sfGFP-TM2β 72-200 recombinant protein.

Proteins were expressed in *E. coli* Rosetta 2 (DE3) pLysS by growing cells to an OD_600_ of 0.4-0.6 in 2xYT (2x yeast extract tryptone) medium at 37 °C, and induced with 0.5 mM IPTG (Isopropyl β-D-1-thiogalactopyranoside) following a cold shock at 4°C. After induction, cells expressing each protein were grown overnight at 16°C. Cells were collected by centrifugation for 20 min at 4,000 rpm, 4°C and resuspended in 20 mM Tris-HCl, pH 8.0, 10 mM imidazole, 2 mM MgCl_2_, 500 mM KCl, 10% (v/v) glycerol, 0.5 mM Tris (2-carboxyethyl) phosphine, and Roche protease inhibitor.

Cells were lysed by sonication, and cell lysate was clarified by centrifugation at 17,000 x g, 4 °C for 0.5 h. The supernatant was bound to Nickel-NTA affinity resin pre-equilibrated with wash buffer (20 mM Tris-HCl, pH 8.0, 500 mM KCl, 30 mM imidazole, 10% (v/v) glycerol, and 0.5 mM Tris(2-carboxyethyl) phosphine) for 1 h at 4 °C. Supernatant was discarded and resin was washed 5 × 30 mL wash buffer (20 mM Tris-HCl, pH 8.0, 500 mM KCl, 30 mM imidazole, 10% (v/v) glycerol, and 0.5 mM Tris(2-carboxyethyl) phosphine). All buffers for the purification of ECOR31 TM2β were additionally supplemented with 1 µM 3’5’-c-di-AMP in the washing and subsequent steps. Protein was eluted in 5 mL elution buffer (20 mM Tris-HCl, pH 8.0, 500 mM KCl, 300 mM imidazole, 10% (v/v) glycerol, and 0.5 mM Tris(2-carboxyethyl) phosphine). Recombinant TEV Protease or SENP2 (SUMO protease 2) with an N-terminal His-tag (BPS Bioscience) was added to the elution for cleavage and dialyzed overnight at 4 °C in a 4,000 MWCO dialysis cassette (Thermo Fisher Scientific) with dialysis buffer (20 mM Tris-HCl, pH 8.0, 500 mM KCl, 30 mM imidazole, 10% (v/v) glycerol, and 0.5 mM Tris(2-carboxyethyl) phosphine). The resultant solution was passed over a 5 mL Ni-NTA Superflow cartridge (Cytiva). Flow-through was collected, concentrated to < 2 mL using a MWCO centrifugal filter (Amicon), and loaded onto a HiLoad 16/600 Superdex 200 pg column (Cytiva). Elution was isocratic (20 mM Tris-HCl, pH 8.0, 500 mM KCl, 10%, 1 mM TCEP) glycerol monitored by A280, peaks were pooled, concentrated, snap-frozen, and stored at -80 °C.

### Analysis of recombinant protein

Purified protein was analyzed by SDS-PAGE. Samples were prepared in 1X Protein Loading Dye (50 mM Tris-HCl, pH 6.8, 15 mM EDTA, 6% (v/v) glycerol, 10% SDS, and bromophenol blue), heated at 95 °C for 3 min, loaded onto a 12% Mini-Protean TGX Precast Protein Gel (Bio-Rad) and run at 125 V until the dye front reached the bottom of the gel. Gels were stained in 30% ethanol, 10% glacial acetic acid in water, 1g R-250 Coomassie and destained in 40% ethanol, 10% glacial acetic acid in water.

### Crystallization and structure determination

Crystals of mCpol in its apo form or bound to a hydrolysis-resistant amine-modified analog of c-di-AMP (ApNHpp, Sapphire North America, NU-449-1) were grown at 20°C using the hanging drop vapor diffusion method. A 1 µL solution of 3 mg/mL mCpol in 20 mM Tris-HCl, pH 8.0, 500 mM NaCl, 5% glycerol was mixed with 1 µL 0.1 M Bicine pH 8.0, 15% PEG 1500 and supplemented with 0.2 µL 10 mM MgCl_2_ with 10 mM of ApNHpp. Single crystals appeared within 3 days and were cryoprotected in a solution of mother liquor with 30% glycerol before being flash cooled in liquid nitrogen.

Data for mCpol bound to ApNHpp were collected *via* fine-phi slicing using 0.2° oscillations at beamline 12-2 at Stanford Synchrotron Radiation Lightsource at SLAC National Accelerator Laboratory. X-ray diffraction data were measured to 2.28 Å resolution.

### Processing and refinement of crystallographic data

Crystallographic data were processed with the SSRL autoxds script with an I/sigI cutoff ≥ 1.50. Crystals displayed moderate anisotropic X-ray diffraction with some diffraction extending beyond 2.0 Å resolution. However, the resolution was isotropically truncated to 2.28 Å resolution to generate a robust complete dataset. The structure was solved by molecular replacement using a ColabFold-generated^48^ model (pLDDT = 95.12) of mCpol with residual residues from the C-terminal cleavage site (mCpol-ENLYFQ) in PHENIX^49^. Molecular replacement successfully identified the placement of two mCpol monomers as indicated by the log-likelihood gain (LLG) 556.03 and the translation-function Z-score (TFZ) 23.6. The structure was refined using PHENIX ^50^ including simulated annealing, non-crystallographic symmetry, and TLS parameters. In each protein monomer (chains A and B) residues 121-125 were disordered and thus not included in the model. The model was built and adjusted using COOT^51^. The structure was refined to a final R_free_ and R_work_ of 24.88% and 22.43% respectively (Supplementary Table 1). Supplementary Table 1 shows data processing and model refinement statistics. Atomic coordinates and structures factors have been deposited to the Protein Data Bank.

### Sequence alignment of mCpol domains

Proteins in the Pfam entry PF18182 “minimal CRISPR polymerase domain” were downloaded and used to create an alignment of 163 mCpol sequences. Sequences were trimmed and aligned using MAFFT alignment with default parameters in Geneious Prime v2023.2.1

### Thin layer chromatography (TLC) of cyclase products

Recombinant enzymes were assessed for cyclase activity by *in vitro* reactions with nucleoside triphosphates and analysis by TLC. Cyclase activity assays were initiated by the addition of recombinant enzyme (40 µM final) in reaction buffer (50 mM Tris, pH 8.0, 10 mM MgCl2, 100 mM NaCl) to 5 mM NTP (Thermo Scientific). The reaction mixture was incubated at 37°C for 18 h and stopped by vortexing for 20 s.

Recombinant enzymes were assessed for cyclic dinucleotide (c-di-AMP) degradation in biochemical reactions containing buffered c-di-AMP and products were analyzed by TLC. Reactions were initiated by the addition of recombinant enzyme (40 µM) in reaction buffer (50 mM Tris, pH 8.0, 10 mM MgCl2, 100 mM NaCl) to 1.25 mM c-di-AMP (Biolog or MedChemExpress). The reaction mixture was incubated at 37 °C from 10 min to 18 h and stopped by vortexing for 20 s.

To silica gel 60 matrix TLC plates (5 cm x 10 cm) on glass support with fluorescent indicator 254 (Supelco) was added 2 µL in vitro enzymatic reaction mix. Separation was performed in an eluent of *n*-propanol/ammonium hydroxide/water (11:7:2 v/v/v) for 45 min. The plate was allowed to dry fully and visualized with a short-wave ultraviolet light source at 254 nm.

### S1 nuclease digestion of cyclase products

To determine the linkage identity of products formed in vitro, cyclase reactions described previously were subjected to digestion by S1 nuclease. To a reaction containing 1X S1 nuclease buffer (40 mM sodium acetate (pH 4.5 at 25 °C), 300 mM NaCl_2_) was added 20 µL vortex-inactivated cyclase reaction and 200 U S1 nuclease (Thermo Scientific) to a total volume of 40 µL. The reaction was incubated at 37°C for 4 h and inactivated by the addition of 2 µL of 0.5 M EDTA with heating (70°C for 10 min).

### Preparation of extracts for mass spectrometry

mCpol-ATP reactions were diluted 1:2 in deionized H_2_O, centrifuged at 13,000 x g for 15 min and used directly in subsequent analysis. Species identified by TLC were directly analyzed by TLC-MS. Silica containing the product was scraped away from the TLC plate and added to 40 µL water. The resultant slurry was vortexed and heated at 30°C for 10 min and then centrifuged at 13,000 x g for 15 min. The supernatant was removed for subsequent analysis.

### Cyclase product analysis by HPLC-MS

Cyclase product extracts were analyzed using a liquid chromatography (LC) system (1200 series, Agilent Technologies, Santa Clara, CA) that was connected in line with an LTQ-Orbitrap-XL mass spectrometer equipped with an electrospray ionization (ESI) source (Thermo Fisher Scientific, Waltham, MA). The LC system was equipped with a G1322A solvent degasser, G1311A quaternary pump, G1316A thermostatted column compartment, and G1329A autosampler unit (Agilent). The column compartment was equipped with an Ultra C18 column (length: 150 mm, inner diameter: 2.1 mm, particle size: 3 µm, catalog number: 9174362, Restek, Bellefonte, PA). Ammonium acetate (≥98%, Sigma-Aldrich, St. Louis, MO), methanol (Optima LC-MS grade, 99.9% minimum, Fisher, Pittsburgh, PA) and water purified to a resistivity of 18.2 MΩ·cm (at 25 °C) using a Milli-Q Gradient ultrapure water purification system (Millipore, Billerica, MA) were used to prepare mobile phase solvents. Mobile phase solvent A was water and mobile phase solvent B was methanol, both of which contained 10 mM ammonium acetate. The elution program consisted of isocratic flow at 0.5% (volume/volume) B for 2 min, a linear gradient to 99.5% B over 2 min, isocratic flow at 99.5% B for 4 min, a linear gradient to 0.5% B over 1 min, and isocratic flow at 0.5% B for 21 min, at a flow rate of 100 µL/min. The column compartment was maintained at 30 °C and the sample injection volume was 1 µL. External mass calibration was performed in the positive ion mode using the Pierce LTQ ESI positive ion calibration solution (catalog number 88322, Thermo Fisher Scientific). Full-scan, high-resolution mass spectra were acquired in the positive ion mode over the range of mass-to-charge ratio (*m*/*z*) = 300 to 2000, using the Orbitrap mass analyzer, in profile format, with a mass resolution setting of 60,000 (at *m*/*z* = 400, measured at full width at half-maximum peak height, FWHM). Tandem mass (MS/MS or MS^2^) spectra were acquired using collision-induced dissociation (CID) in the linear ion trap, in centroid format, with the following parameters: isolation width = 3 *m*/*z* units, normalized collision energy = 28%, activation Q = 0.25, and activation time = 30 ms. Data acquisition was controlled using Xcalibur software (version 2.0.7, Thermo Fisher Scientific).

### Mass spectrometry data processing

Raw data were converted to mzXML format using msconvert 3.0.19052.1 from the Galaxy platform^52^. Data were then processed using the open source software MZmine 3.9.0. Compound identification was performed by differential mass spectrometry of in vitro reactions with active and inactive enzymes, and by comparing retention times and mass-to-charge ratios (m/z) with those of chemical standards.

### Gel shift assays

Reactions were performed using purified 2TMβ Δ2TM (residues 72-200) and phosphodiesterase (PDE) enzymes by adding PDE (pigeon pox virus PDE wild-type and H72A/H167R^42^) at a final concentration of 5 mg/mL to a solution containing 0.3 mg/mL of His-GFP-SUMO-2TMβ in 1X c-di-AMP-containing storage buffer (20 mM Tris-HCl, pH 8.0, 500 mM KCl, 10%, 1 mM TCEP, and 1 µM 3’5’-cyclic di-AMP) and incubated for 10 min at 37 °C, then transferred to ice. On ice, native PAGE loading buffer (50 mM Tris-Cl, 0.1% bromophenol blue, 10% glycerol, 100 mM DTT) was added 1:1 to reactions, applied to an 8% native PAGE gel at 4°C, and electrophoresed for 1 h 45 min at 150 V. The gel was imaged using a BioRad ChemiDoc on the SYBR Safe setting.

### Fluorescence microscopy

Inducing agents or water were added to 400 µL of log phase cultures at OD_600_ 0.12-0.2 (final concentrations: aTc = 200 nM, CV = 50 nM), which were then incubated in a roller for 50 min at 30°C. 100 µL of stain mix (LB, DAPI and MitoTracker Green FM) were added to the cultures for a final stain concentration of 4 µg/mL DAPI and 2.5 µg/mL MitoTracker Green FM and incubated, rolling, for an additional 10 min at 30°C. 20 µL of culture were spotted on an agarose imaging pad (25% LB, 1% agarose) in single-well concavity slides for imaging. Cells were imaged at room temperature with eight 0.2 µm slices in the Z-axis. Images were collected on a DeltaVision Elite Deconvolution microscope (Applied Precision, Issaquah, WA, USA) using DeltaVision SoftWoRx (version 6.5.2) and figure panels were prepared in Adobe Photoshop (21.2.0) and Adobe Illustrator (24.2). Imaging was performed in biological duplicate.

### Sequence mining and genomic neighborhood analysis of mCpol

A collection of mCpol encoding genes was curated using the following strategy. First, a DALI-search was performed against representatives of the clustered AlphaFold Database (AFDB) using an AlphaFold2 model of the mCpol (A0A426EXS8). After filtering for a Z-score of 12, a structurally informed multiple sequence alignment (MSA) was generated and used for phylogenetic analysis as described previously^53^. The clade containing mCpol representatives (N=5) was extracted from the resulting tree, and the selection was then expanded to include all cluster members (N=29). A structurally informed MSA was generated from this set, and used as input for an HMM-search against NCBI_NR_Aug_2018 database on the MPI Bioinformatics Toolkit website (MSA enrichment iterations using HHblits: 1, E-value cutoff for reporting: e-10, Max target hits: 1000). This set of mCpol candidates (N=592) was aligned using MAFFT v7.490 in Geneious Prime 2023.0.1 (Algorithm: E-INS-i, Scoring matrix: BLOSUM62, Gap open penalty: 1.3). Fragmented mCpol genes were removed from the alignment, and the remaining genes were used for neighborhood analysis (N=593). A custom Python script was used to retrieve Genbank files of mCpol-containing loci. Only contigs longer than 20kb were kept for further analysis (N=505). The resultings contigs were converted to FASTA format and deduplicated using mmseqs^54^ easy-cluster (−c 0.5 --min-seq-id 0.5 --dbtype 2 -- cluster-mode 2). The final set of sequences (N=252) were then subjected to defense association analysis via DefenseFinder Web Service (https://defensefinder.mdmlab.fr/) ^35,55^. The outputs of DefenseFinder were then further processed using a custom Python script to add defense system hit annotations to the Genbank files. This final output was used to investigate co-occurrence of mCpol with CBASS and other types of defense systems.

### Taxonomic analysis of mCpol-encoding genes

To determine the taxonomic distribution of mCpol homologs, we retrieved the NCBI Taxonomic Identifier (taxID) associated with each protein sequence. Protein IDs were supplied to a custom script that queries the NCBI Entrez system via E-utilities (esearch, elink, efetch, and xtract)^56^ to extract both the taxID. For protein IDs that failed automated retrieval, taxonomic information was manually obtained using NCBI’s web interface. For each taxID, we used a separate custom Python script that calls the NCBI Datasets^57^ command-line tool to obtain taxonomy summaries in JSON format and subsequently parses the taxonomic ranks saved as a tsv. A Jupyter notebook was used to further cross-referenced protein IDs with locus tag information from associated annotation files. Merging and reformatting of the dataset was performed in Python using pandas, yielding a finalized tab-delimited table containing accession IDs, taxonomic lineages, and protein identifiers for downstream analysis.

