## Supplementary Material for "A miniature CRISPR-Cas10 enzyme confers immunity by an inverse signaling pathway"

**Supplementary Figures**

**Supplementary Fig. 1. Representative plaque assays for phage sensitivity to MISS.**
**Supplementary Fig. 2. Sequence alignment of 163 mCpol representatives from PF18182.**
**Supplementary Fig. 3. Thin Layer Chromatography-Mass Spectrometry (TLC-MS) characterization of products formed when recombinant mCpol is added to ATP.**
**Supplementary Fig. 4. Purification of ECOR31 for crystallization.**
**Supplementary Fig. 5. mCpol crystal structure reveals mechanism of adenine base specificity.**
**Supplementary Fig. 6. Increasing expression of MISS yields decreasing levels of phage defense.**
**Supplementary Fig. 7. Conservation of β-barrell domain nucleotide binding residues.**
**Supplementary Fig. 8. Genetic contributions towards phage resistance of MISS.**
**Supplementary Fig. 9. Representative plaque assays for CRISPRi-ART repression of phage encoded MISS activator proteins.**
**Supplementary Fig. 10. Mutant controls for MISS Activator assays.**
**Supplementary Fig. 11. Phylogenetic analysis of mCpol proteins and representative MISS loci.**

**Supplementary Table 1. X-ray data collection and refinement statistics**

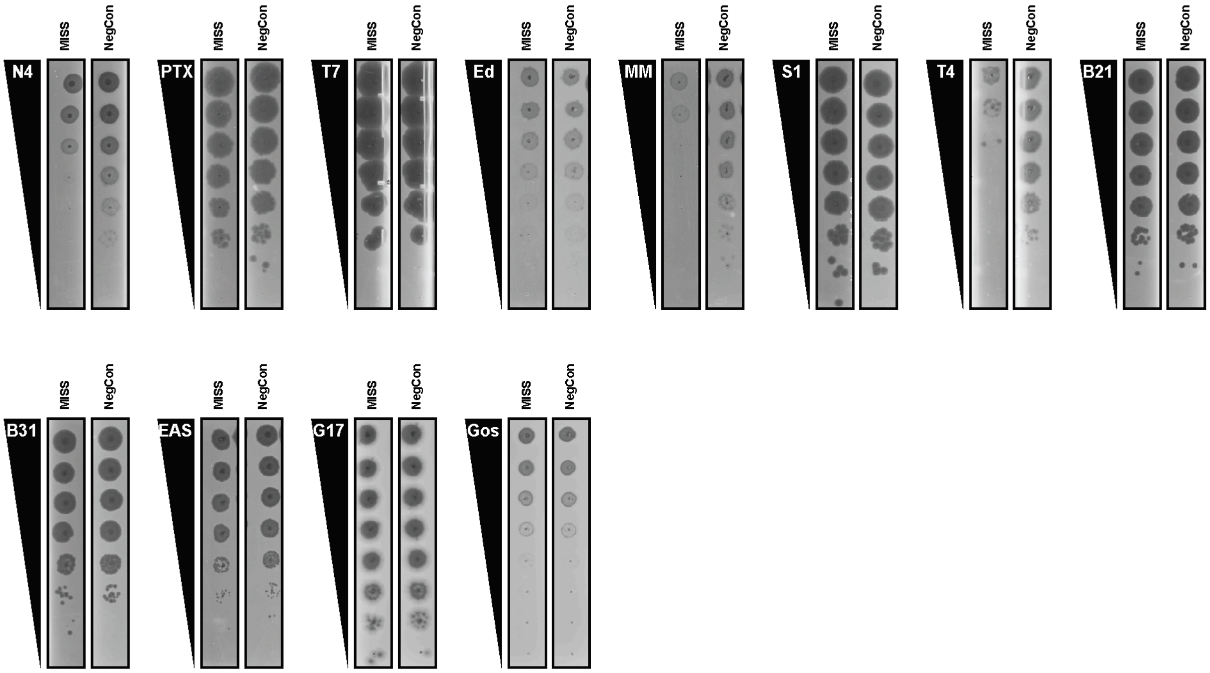


**Supplementary Fig. 1. Representative plaque assays for phage sensitivity to MISS.** Plaque assays of phages shown in the order they appear in Figure 1 and their sensitivity to MISS-mediated antiphage defense. Plaque assays shown are representative images from 3 biological replicates.

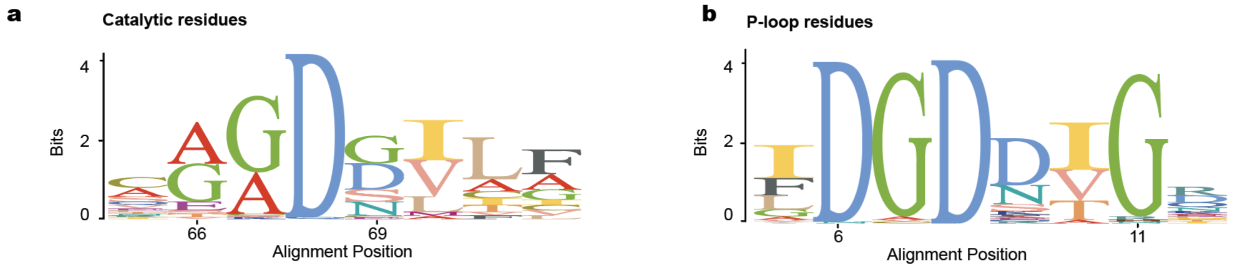


**Supplementary Fig. 2. Sequence alignment of 163 mCpol representatives from PF18182.** (a) Semi-conserved GGDD motif (AADG in ECOR31 mCpol). (b) Conserved active site residues.

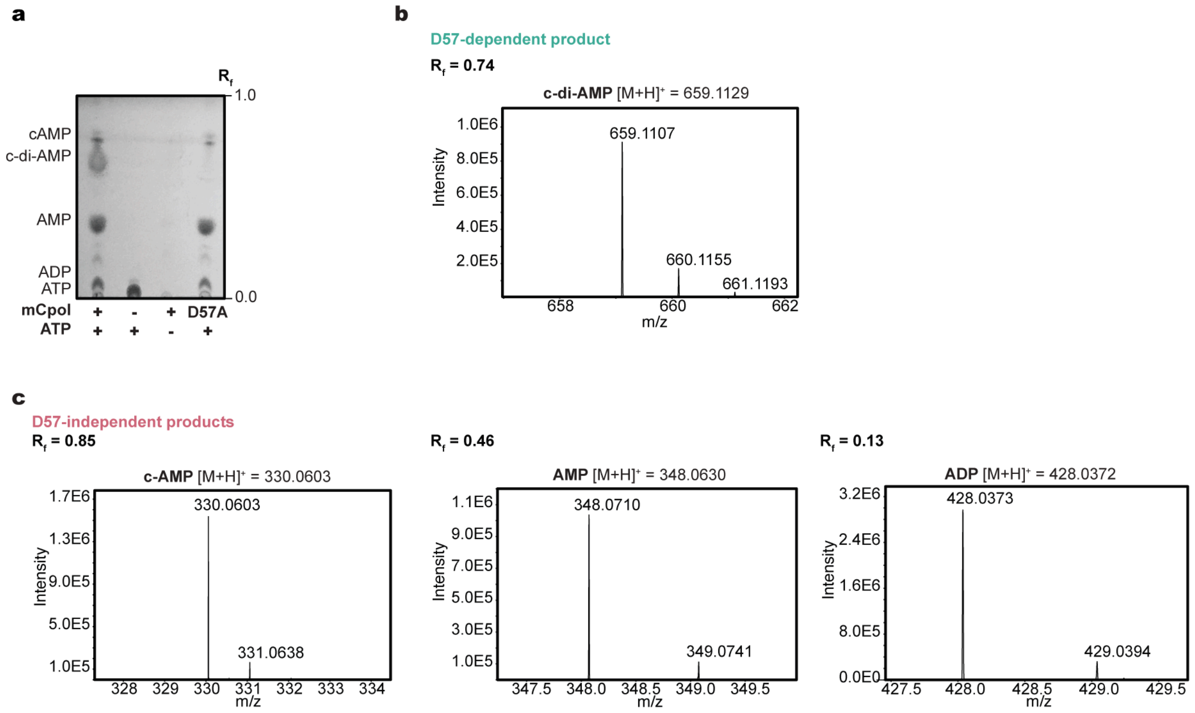


**Supplementary Fig. 3. Thin Layer Chromatography-Mass Spectrometry (TLC-MS) characterization of products formed when recombinant mCpol is added to ATP.**

**(a)** A representative TLC plate showing reaction between mCpol or mCpol D57A and ATP by $R_f$.

**(b)** MS of the mCpol D57-dependent product spot.

**(c)** MS of the 3 major product spots that are not dependent upon the conserved aspartate in mCpol.

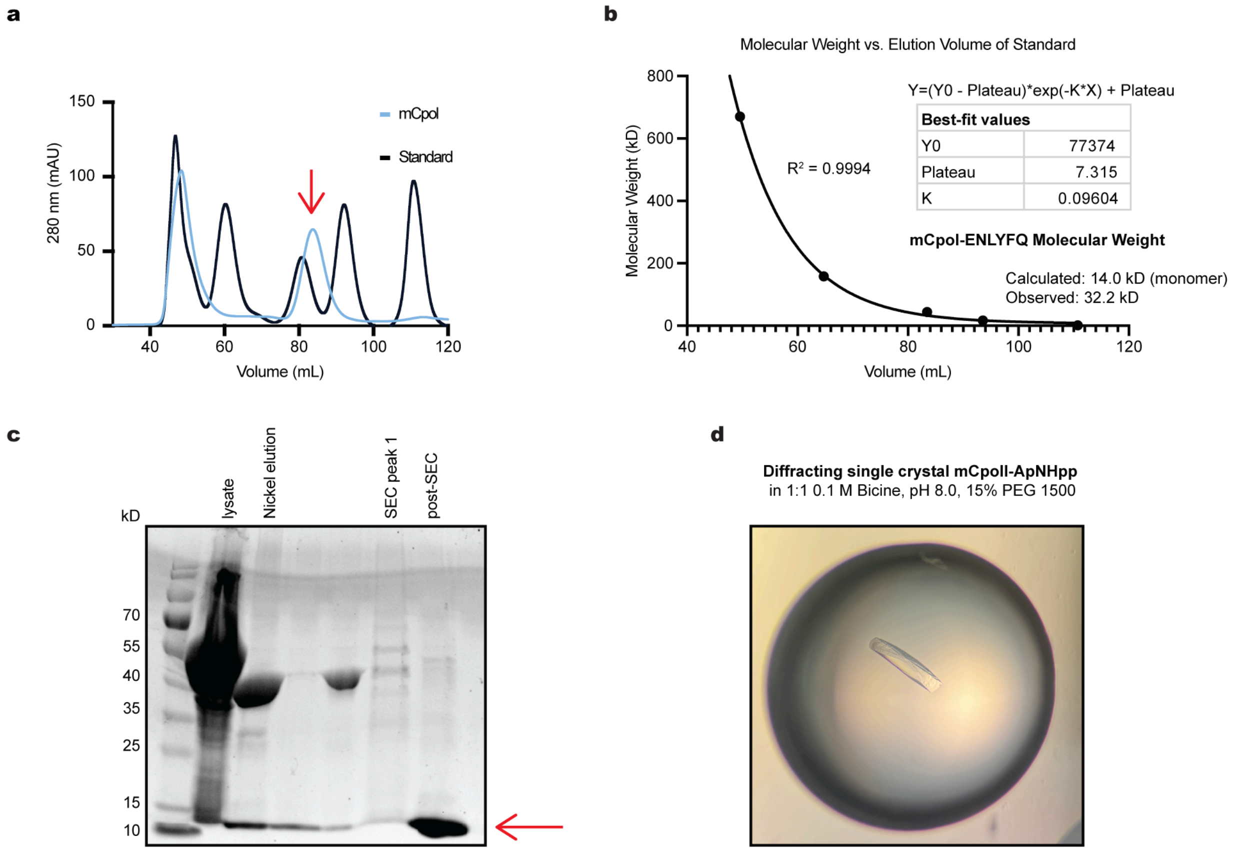


**Supplementary Fig. 4. Purification of ECOR31 mCpol for crystallization.**

**(a)** Size exclusion chromatography (SEC) of ECOR31 mCpol with ENLYFQ cleavage site (red arrow) versus a molecular weight standard.

**(b)** Determination of mCpol multimerization state by comparison with a molecular weight standard.

**(c)** Purification of mCpol for crystallography (trays set up with post-SEC fraction, red arrow).

**(d)** Single crystal resulting from the 0.1 M bicine, pH 8.0, 15% PEG 1500 condition, mixed 1:1 with protein and supplemented with 0.2 equivalents of 10 mM of ApNHp in 10 mM $MgCl_2$.

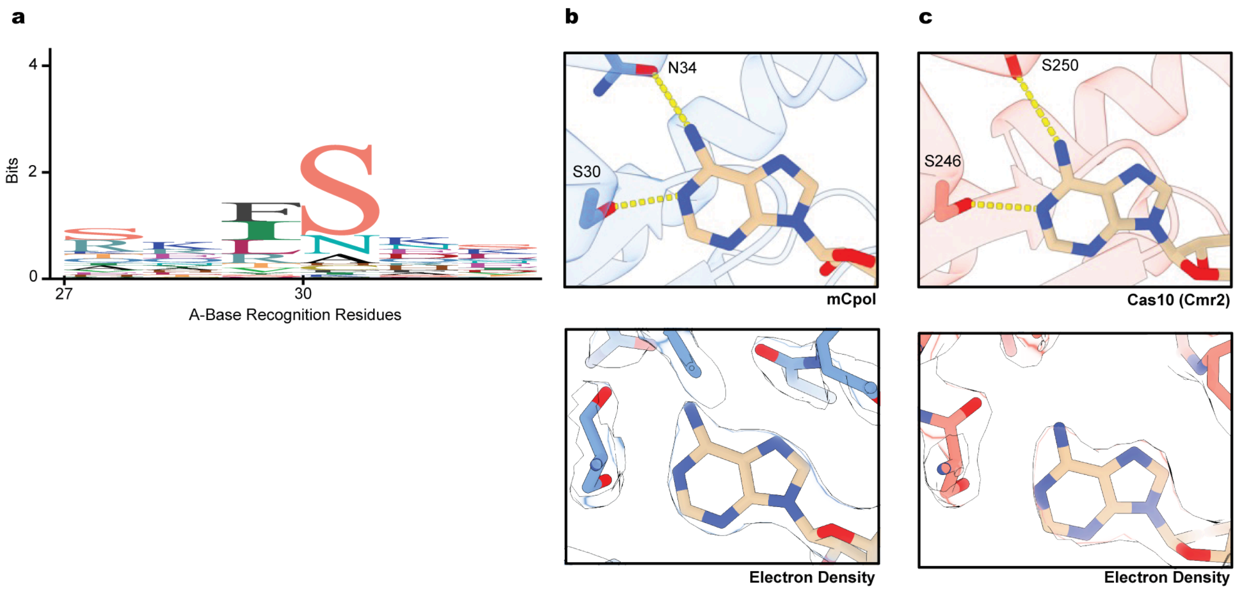


**Supplementary Fig. 5. mCpol crystal structure reveals mechanism of adenine base specificity.**

**(a)** Conservation of S30 in an alignment of 163 mCpol domain-containing proteins.

**(b)** Base specific contacts made between mCpol and the adenine base and corresponding electron density.

**(c)** Base specific contacts made between Cas10 (Cmr2, PDB ID: 3w2w) and the adenine base and corresponding electron density.

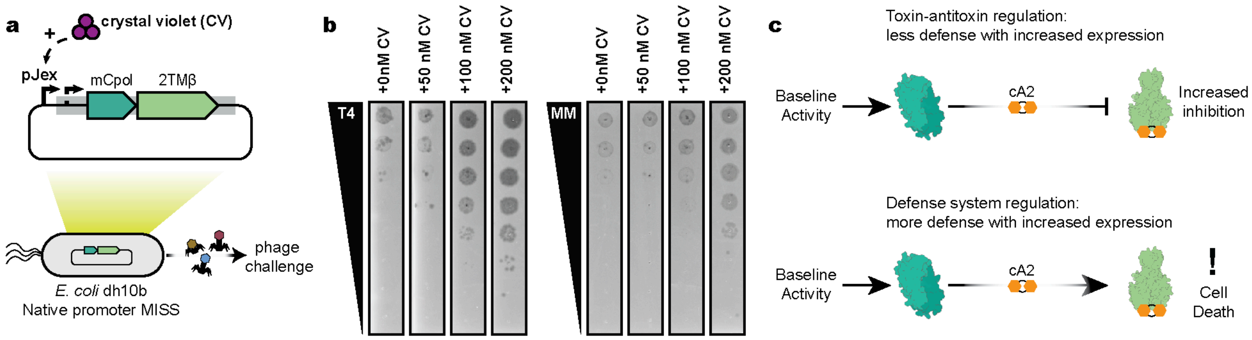


**Supplementary Fig. 6. Increasing expression of MISS yields decreasing levels of phage defense.** (a) Overview of phage infection experiment with additional mCpol induction. (b) Plaque assays of T4 and MM02 phages in the presence of increasing expression levels of MISS at specified concentrations of CV. Plaque assays are representative images of 3 biological replicates. (c) Comparison of toxin-antitoxin (top) and defense system architectures (bottom). Increased expression of a toxin-antitoxin signaling system would also yield increased inhibition of the toxin and less defense. Increased expression of a defense system would yield higher levels of defense and potentially incidental cell death.

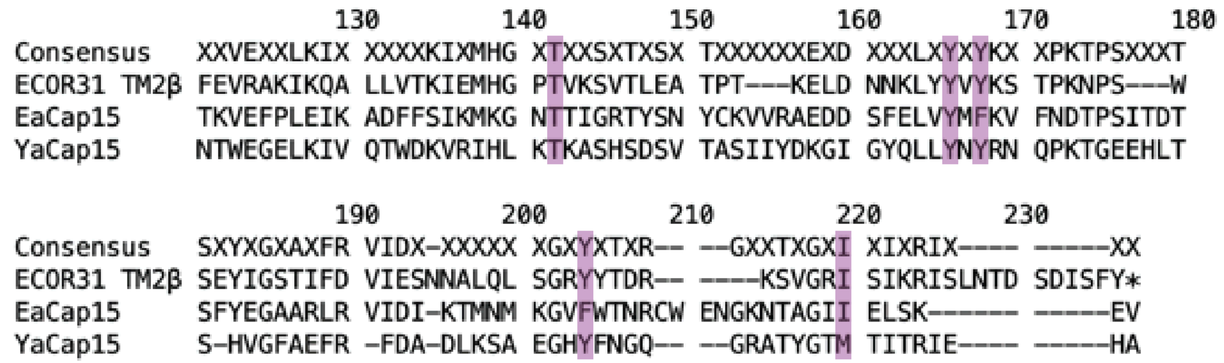


**Supplementary Fig. 7. Conservation of β-barrel domain nucleotide binding residues.** Alignment of a segment of the β-barrell nucleotide binding domain of *Escherichia albertii* Cap15 and *Yersinia* Cap 15 with 2TMβ from ECOR31 MISS. Highlighted residues (corresponding to T129, Y153, Y155, Y188 and M200 in *Ea*Cap15) were found to be functionally important for cyclic dinucleotide binding by Duncan-Lowey et al.[17]

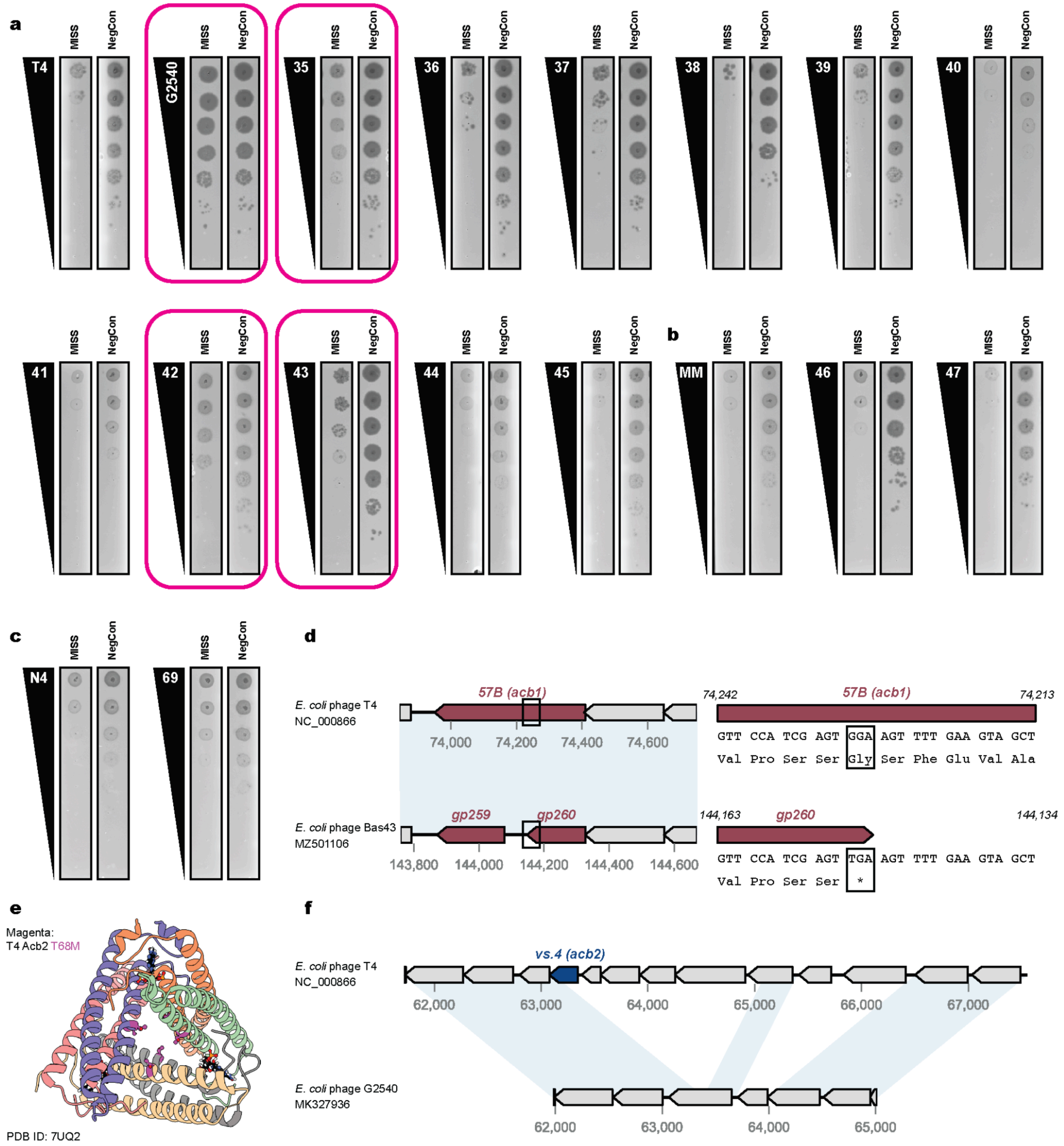


**Supplementary Figure 8. Genetic contributions towards phage resistance of MISS.**

(a) Plaque assays for T4-like phages from DSMZ (G2540) or the BASEL collection (Bas35-45). Resistant phenotypes are highlighted in pink.
(b) Plaque assays for MM02-like phages from the BASEL collection (Bas46-47).
(c) Plaque assays for N4-like phages from the BASEL collection (Bas69). For panels a-c, plaque assays are representative of 3 biological replicates.
(d) Comparative genomics highlights a nonsense mutation at the Bas43 *gp260* locus (a T4 Acb1 homolog). Regions of high nucleotide similarity as identified by ProgressiveMauve [58] are

highlighted.
(e) Mutation decreasing T4 Acb2 binding affinity, T68M [40], in Bas35 shown in magenta overlaid on PDB 7UQ2.
(f) Comparative genomics highlights a multigene deletion at the *acb2* locus in between T4 and G2540, including complete loss of *acb2*. Regions of high nucleotide similarity as identified by ProgressiveMauve [58] are highlighted.

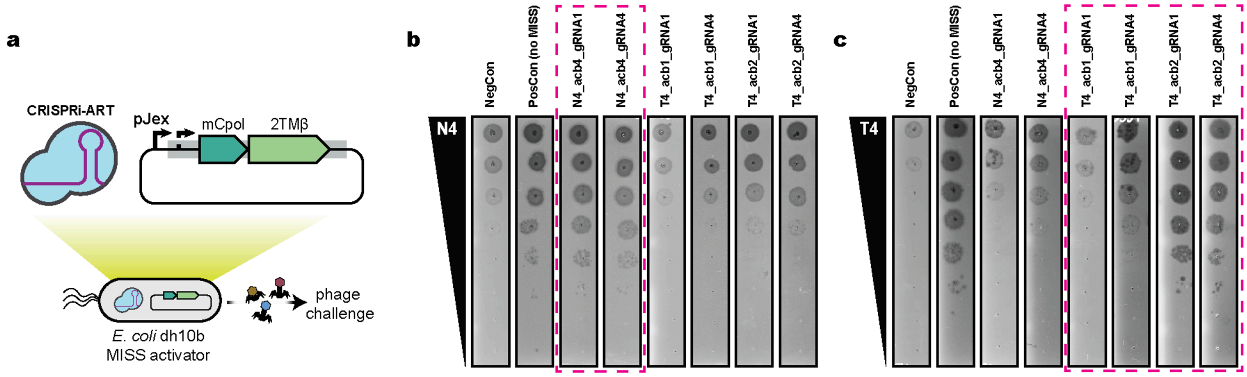


**Supplementary Fig. 9. Representative plaque assays for CRISPRi-ART repression of phage encoded MISS activator proteins.** (a) CRISPRi-ART screening of phage encoded Acb proteins to restore phage infectivity in the context of MISS. For all experiments, CRISPRi-ART was expressed at +20nM aTc. NegCon represents a CRISPRi-ART crRNA targeting a transcript not present in the experiment. PosCon represents a non-targeting CRISPRi-ART crRNA in the absence of MISS. CRISPRi-ART conditions targeting a phage-encoded Acb protein in the target phage are bounded in dashed magenta lines. All plaque assays shown are representative images of 3 biological replicates. (b) N4 infection for CRISPRi-ART targeting of Acb proteins in the presence of MISS. (c) T4 infection for CRISPRi-ART targeting of Acb proteins in the presence of MISS

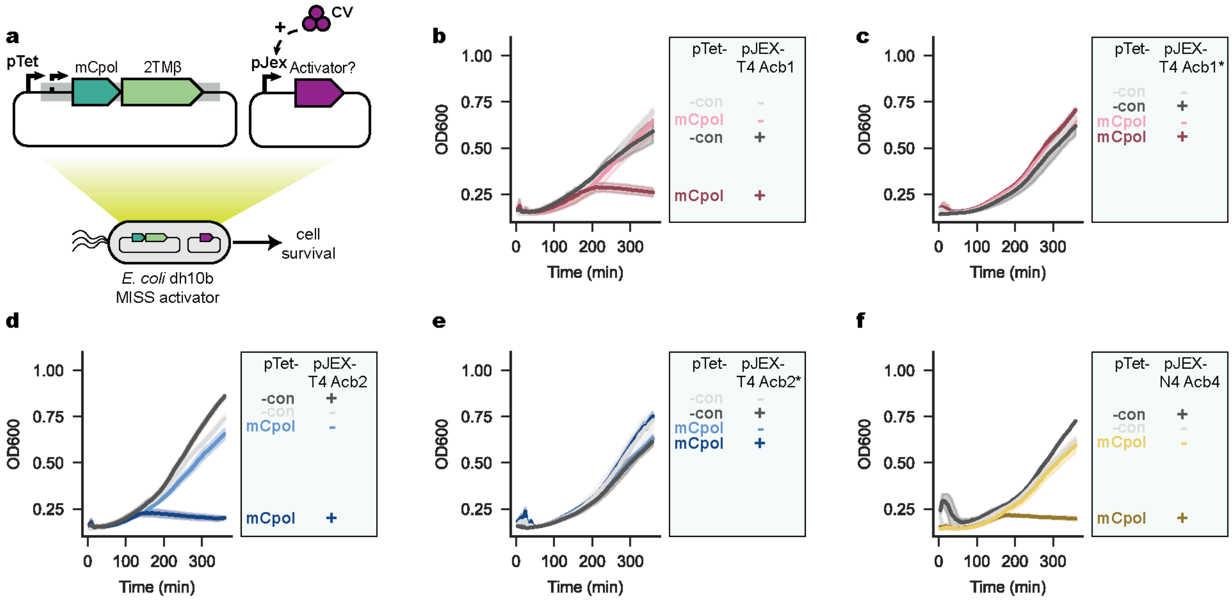


**Supplementary Fig. 10. Mutant controls for MISS Activator assays.** (a) Overview of activator assays in the context of MISS under control of its native promoter. For activator assays in panels b-f candidate MISS activator proteins are expressed at +0 or +125nM CV. (b) T4 Acb1 expression is sufficient to activate MISS toxicity. (c) Catalytically deactivated mutant of T4 Acb1 () expressed is insufficient to activate MISS toxicity. (d) T4 Acb2 expression is sufficient to activate MISS toxicity. (e) Binding-deficient mutant of T4 Acb2 (Y8A) expression is insufficient to activate MISS toxicity. (f) N4 Acb4 expression is sufficient to activate MISS toxicity. Data from Figure 4d, 4e, 4f are repeated here for comparative purposes.

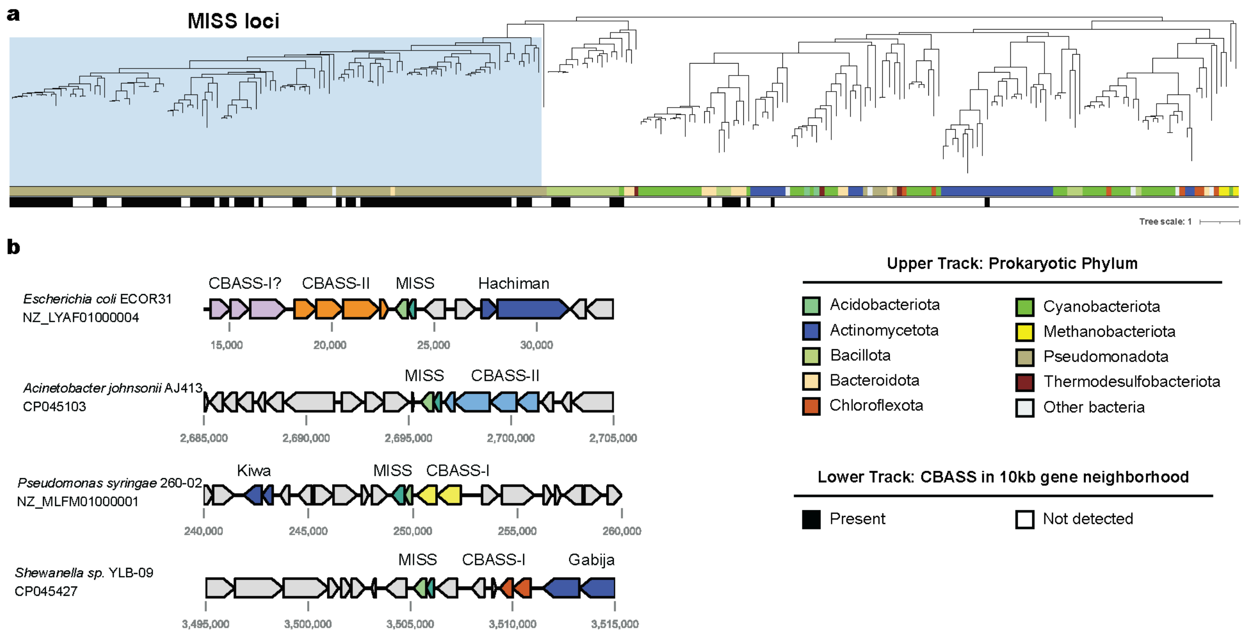


**Supplementary Fig. 11. Phylogenetic analysis of mCpol proteins and representative MISS loci.** (a) Structure-guided phylogenetic tree for the mCpol protein domain. Top track represents host phylum. Bottom track represents CBASS association in a 10kb neighborhood of the mCpol-containing protein. MISS clade investigated in this study is highlighted in light blue. (b) Representative MISS loci highlighting distinct CBASS systems and other defense systems in the 10kb gene neighborhood.

**Supplementary Table 1. X-ray data collection and refinement statistics**

| PDB ID | | mCpol-ApNHpp<br>9NWN |
|---|---|---|
| **Data collection[a,b]** | | |
| Space group | | $P2_12_12$ |
| Cell Dimensions | | |
| | *a, b, c* (Å) | 49.1, 56.8, 93.7 |
| | α, β, γ (°) | 90, 90, 90 |
| Resolution (Å) | | 48.55-2.28 (2.37-2.28) |
| $R_{merge}$ (%) | | 14.6 (75.9) |
| $R_{pim}$ (%) | | 6.1 (31.1) |
| $CC_{1/2}$ (%) | | 99.8 (88.4) |
| <I/σI> | | 6.8(2.0) |
| Completeness (%) | | 98.8 (100.0) |
| Redundancy | | 6.6 (6.9) |
| Wilson *B*-factor (Å$^2$) | | 35.28 |
| **Refinement and Validation** | | |
| Resolution (Å) | | 48.55-2.28 (2.37-2.28) |
| Unique Reflections | | 12,282 (1,349) |
| Number of atoms | | |
| | Protein | 1,897 |
| | Ligand | 64 |
| $R_{work}/R_{free}$ (%) | | 22.4/24.9 |
| R.m.s. deviations | | |
| | Bond lengths (Å) | 0.014 |
| | Bond angles (˚) | 1.46 |
| Poor rotamers (%) | | 2.35 |
| Ramachandran plot | | |
| | Favored (%) | 94.5 |
| | Allowed (%) | 5.51 |
| | Disallowed (%) | 0 |
| Average *B*-factor (Å$^2$) | | 48.86 |

[a]For each structure reported, data were derived from a single crystal.
[b]Numbers in parentheses correspond to the highest resolution shell
